# A study of psychological pain in substance use disorder and its relationship to treatment outcome

**DOI:** 10.1101/613737

**Authors:** Steven Mee, Blynn G Bunney, Ken Fujimoto, John Penner, Garrett Seward, William E Bunney, Christopher Reist

## Abstract

Substance Use Disorder (SUD) is a major public health concern affecting an estimated 22.5 million individuals in the United States. The primary aim of this study was to characterize psychological pain in a cohort of patients participating in outpatient substance abuse treatment. A secondary aim was to determine the relationships between pre-treatment assessments of psychological pain, depression, anxiety and hopelessness with treatment retention time and completion rates. Data was analyzed from 289 patients enrolled in an outpatient community drug treatment clinic that provides mental healthcare to the underserved. A previously determined threshold score on the Mee-Bunney Psychological Pain Assessment Scale (MBP) was utilized to group patients into high and low-moderate scoring subgroups. The higher pain group reported increased levels of anxiety, hopelessness and depression compared to those in the low-moderate pain group. Additionally, patients scoring in the higher psychological pain group exhibited reduced retention times in treatment and more than two-fold increased odds of dropout relative to patients with lower pre-treatment levels of psychological pain. Among all assessments, the correlation between psychological pain and treatment retention time was strongest. To our knowledge, this is the first study to demonstrate that psychological pain is an important construct that correlates with relevant clinical outcomes in substance abuse treatment. Further, pre-treatment screening for psychological pain may be of benefit in identifying higher-risk patients in need of targeted additional clinical resources to improve treatment retention and completion rates.

## INTRODUCTION

Substance Use Disorder (SUD) is a major public health concern affecting an estimated 22.5 million individuals in the United States (1). In 2017, 70,237 deaths were attributed to drug overdoses—a significant increase of 9.6% over the past year (2). Nationally, substance-related addiction incurs a financial burden exceeding $400 billion per year including expenses related to lost work productivity, healthcare and drug-related crime (3). There are 14,500 drug treatment centers in the US but only a relatively small number of individuals (11%) enter treatment despite the fact that many programs are supported by local, State and Federal government funding (4). Drug treatment programs continue to strive for improving program outcomes, however, data shows that they are maximally effective when patients remain in treatment (length of stay-LOS) for an average of 90 days or more (5, 6). Program completion is associated with better health, fewer readmissions, less criminal activity (7) and lower mortality rates (8).

Psychological pain is emerging as an important construct based on data from our studies (9,10) and others (11–19). This form of pain is also described as mental pain, emotional pain, social pain and psychache (20–22). There is very little data, however, on characterizing psychological pain in addiction and in addiction treatment populations. Use of addictive substances has been viewed as a strategy to suppress negative emotions so that the control of mental pain (e.g., drug seeking induced by acute stress) is the objective rather than the pleasure-seeking associated with substances of abuse (23). Within this framework, it might be expected that the greater the level of psychological pain, the more serious the addiction. A study in Portugal that assessed psychological pain in an addicted population using a translated (English to Portuguese) abbreviated version (24 items) of the Orbach & Mikulincer Mental Pain Scale (OMMP) found a small to moderate but positive correlation between mental pain and the severity of addiction (24).

Poor retention rates (inadequate LOS) and failure to complete (‘Dropout’) substance treatment programs pose major clinical challenges (25) to successful treatment. Of those SUD individuals entering a program complete treatment depending on the type of abused substance and whether therapy is offered as in- or outpatient (7). Programs often have no predetermined time period for completion as relapse rates are high and it is common for patients to briefly leave then re-enter treatment. In the case of dropouts, however, patients often terminate treatment early without returning. Studies predicting individuals at high risk for non-completion show that demographics, alone, are relatively poor indicators of dropout risk (7). An adequate length of stay (LOS) is considered an important influential factor for completion whereas early dropout from treatment is associated with outcomes comparable to no treatment (26).

A Norwegian study addressing ‘mental distress’ and variables related to dropout included 454 patients from five inpatient SUD centers used a brief version (10-item) of the Hopkins Symptom Checklist (HSCL). The HSCL assesses obsessive-compulsivity, somatization, anxiety and depression. Interestingly a high score on the HSCL, which the investigators interpreted as ‘mental distress’, was associated with treatment dropout. The HSCL, however, does not specifically define or assess psychological pain. Psychiatric diagnoses including mood disorder, anxiety, PTSD and personality disorder did not significantly differentiate dropouts from completers (27).

This study was undertaken to characterize psychological pain using the Mee-Bunney psychological pain scale (MBP) (9, 10) in a substance dependent population. We hypothesized that the construct of psychological pain could be reliably assessed in this population and would correlate with measures of depression and anxiety as previously observed in major depressive episodes and patients with suicidality (9, 10). In addition, to build on these findings, we tested the utility of a previously established threshold for high psychological pain as an indicator of risk for negative treatment outcomes. Our earlier data also showed that patients scoring high for psychological pain upon intake also had higher acuity on other clinical assessments. The MBP is a brief 10-item self-report instrument designed for use in a variety of clinical settings. Items are rated on a 5-point Likert Scale and include categories such as current and past (within 3 months) psychological pain levels, intensity, and tolerance (e.g., how much psychological pain can you tolerate before it becomes unbearable?). We previously documented higher levels of psychological pain in patients with major depressive episodes (MDEs) compared to healthy controls (10) where a secondary finding included a significant correlation between psychological pain and suicidality scores obtained from the Suicide Behavior Questionnaire (28). In a follow-up study we examined psychological pain as a pre-treatment risk indicator for suicidality and serious suicide attempts in U.S. Veterans admitted to a suicide prevention program. Our findings showed that of all the assessments including depression, hopelessness and impulsivity, psychological pain accounted for the most shared variance with suicidality as measured by the Columbia Suicidality Severity Rating Scale (C-SSRS). Using the previously tested MBP threshold score of MBP≥32 [defined as 0.5 SD above the mean of the MDE patients (10)],we identified a subgroup of patients (24/57) scoring high for psychological pain. At a 15-month follow-up, 9 of the 24 patients had a serious suicidal event (as defined by criteria on the C-SSRS). Of those scoring above 32 on the Mee-Bunney scale, 7/9 would have died if not found and one patient completed suicide. Taken together, these results showed that high psychological pain increased the risk for suicidal ideation and serious suicidal events. In addition, they provided preliminary evidence that stratifying patients by psychological pain scores could aid in identifying those experiencing an elevated risk for negative clinical outcomes.

In this study, we hypothesized that psychological pain would correlate with ratings of depression, anxiety and hopelessness consistent with our previous findings in other psychiatric populations. In addition, we tested whether high levels of psychological pain would correlate with an elevated risk for poorer treatment outcomes (i.e., treatment retention times (LOS) and completion rates) compared with lower MBP scoring patients.

## METHODS

### Study design

A retrospective analysis of medical records was conducted for patients enrolled between 2011-2013 in the Substance Abuse Counseling Systems of The Gary Center (La Habra, California (SACS); a community-based outpatient SUD treatment program providing addiction treatment to the underserved. Data collected included demographics, standardized clinical assessments and outcome variables including completion/dropout status and length of stay (LOS). Patients were referred to the SACS program by medical providers, regional non-profit centers, Orange County (OC) courts, legal agencies and the OC Healthcare Agency. In addition, the SACS program was advertised on the internet (http://orange.networkofcare.org/mh/services/agency.aspx?pid=TheGaryCenterSACS_348_2_0). Direct outreach within the Orange County Healthcare Agency was also used to contact clinicians. The SACS clinical program was supported by a grant from the OC Healthcare Agency to provide substance abuse outpatient treatment to all patients, regardless of funding or insurance status. The Institutional Review Board (IRB) of the County of Orange Healthcare Agency approved the study and waived informed consent due to the minimal risk associated with a retrospective chart review. We carefully protected the identity of the patients by assigning each patient a numerical code to ensure privacy. Research personnel conducting chart reviews were blind to the study protocol.

### Subjects

Medical records (N= 529) from January 2011 to December 2013 for male and female patients ≥18 years of age and meeting the DSM-IV criteria for Substance Dependence or Substance Abuse were screened for inclusion in the study. Patients with incomplete medical records or who did not meet admission requirements were excluded from the study so that a total of 289 patient clinical charts were entered into the analyses. Successful program completion was defined as fulfilling all required elements of the clinical program. Data collected in the retrospective chart review included demographics, program length of stay (LOS), completion status and data from clinical rating scales. Detailed socioeconomic variables such as employment, education and marital status were not available. All patients entering the program underwent drug screening at admission and during the course of treatment for alcohol, tetrahydrocannabinol (THC), methamphetamine, cocaine, opiates and benzodiazepines.

### Exclusion criteria

Subjects under age 18 and those who had not agreed to each required random drug screening as well as clinical testing were excluded from the retrospective analyses as were those chart records with missing assessment and/or data relevant to completion status.

### Assessment

Data collected from the intake assessment upon admission included ratings from the Mee-Bunney Psychological Pain Assessment Scale (MBP) (9, 10) the Beck Depression Inventory (BDI)(29), the Beck Hopelessness Scale (30) and the Beck Anxiety Inventory (BAI) (31). Random drug testing conformed to the standards of the Department of Transportation (DOT)-regulated biological fluid testing and included both observed urine and saliva collection.

### Program Completion

was task dependent and determined by successfully completing the core programmatic components as designed by the SACS treatment team. Primary required elements included: attendance in the program >90 days, participation in 24 group sessions (16 process groups and 8 relapse prevention groups); four individual psychotherapy sessions, evidence of weekly attendance at community-based 12-step programs; two psycho-educational classes; and six random, observed drug tests. In order to maximize the opportunity to complete the treatment program and to accommodate relatively brief diversions from treatment (i.e., court hearings and child visitation), there was no predetermined maximum time for completion.

### Program Dropout

(non-Completion status) was defined as not completing the tasks necessary for program completion as described previously and/or non-attendance for greater than 30 days.

### Statistical Methods

Statistical analyses were performed with IBM SPSS software. Multiple regression analyses with predictors entered into the model in blocks were performed to determine correlation coefficients (Pearson) between clinical tests. Group differences were examined using ANOVA, two-tailed t-test (continuous variables) and Chi Square analysis (categorical variables). Significance levels were set at p=0.05. To allow for further examination of outcome variables such as Length of Stay (LOS) and rates of completion or dropout we generated Kaplan-Meier Survival (Retention) curves and performed Log-rank comparison tests with the null hypothesis assuming that the curves would not differ between comparison groups.

## RESULTS

### Demographics

Data from 289 patients (188 males and 71 females) were included in the analyses (Table 1). Patients self-identifying as Hispanic comprised a slight majority of the population (55%). Methamphetamine was the most frequently reported drug of abuse (73.7%) followed by alcohol (64.7%) and cannabis (56.4%). The majority of patients were polysubstance abusers (n=228; 78.9%), while 21.1% (n=61) reported using a single drug of choice. The combined number of drugs used by patients ranged from two (n=116; 40.1%) to five (n=10; 3.5%) with the majority using two substances, methamphetamine and alcohol.

**Table 1.** Demographics

| **Gender/ Age (yrs) ± SD** | **N** | **Percent (%)** |
| --- | --- | --- |
| Hispanic | 159 | 55 |
| Black | 12 | 4.2 |
| Asian | 7 | 2.4 |
| Pacific Islander | 2 | 0.7 |
| Other | 9 | 3.1 |
| **Drug of choice with (+)or without (-) polysubstance** |  |  |
| Alcohol | 205 | 70.9 |
| (+) | 187 | 91.2 |
| (-) | 18 | .09 |
| Tetrahydrocannabinol (THC) | 178 | 61.6 |
| (+) | 163 | 91.6 |
| (-) | 15 | 8.4 |
| Methamphetamine | 238 | 82.4 |
| (+) | 213 | 89.4 |
| (-) | 25 | 11 |
| Cocaine | 68 | 23.5 |
| (+) | 66 | 97.1 |
| (-) | 2 | 2.9 |
| Opiates | 51 | 17.6 |
| (+) | 0 | 17.6 |
| (-) | 51 | 0 |
| Benzodiazepines | 8 | 2.8 |
| (+) | 7 | 87.5 |
| - drug | 1 | 12.5 |
| **Number of drugs abused** |  |  |
| One | 61 | 21.1 |
| Two | 116 | 40.1 |
| Three | 64 | 22.1 |
| Four | 38 | 13.1 |
| Five | 10 | 3.5 |

### Clinical ratings

As described in Table 2 scoring of clinical assessments for all patients indicated low levels of depression (BDI), anxiety (BAI) and hopelessness (BHS). Psychological pain scores were in the low-moderate range based on previous studies in normal and depressed populations (Mee, et al., 2011). Determination of Cronbach’s alpha indicated good internal reliability for all assessment instruments: MBP= .902, BDI=.941, BHS=.876, BAI=.958.

**Table 2:** Mean scores and ratings for psychological pain (MBP), depression (BDI), anxiety (BAI) and hopelessness (BHS)

| Scale | Score (±SD) | Scoring Category |
| --- | --- | --- |
| MBP | 23.57 (8.87) | Low-moderate |
| BAI | 11.98 (14.46) | Mild |
| BHS | 4.8 (4.48) | Mild |
| BDI | 8.54 (6.60) | Minimal |

Significant correlations between the clinical assessments are described in Table 3. The strongest relationship was found between psychological pain (MBP) and depression (BDI).

**Table 3.** Pearson correlation coefficients between rating scales

|  | MBP | BDI | BHS | BAI |
| --- | --- | --- | --- | --- |
| MBP<sup>a</sup> | 1.00 |  |  |  |
| BDI<sup>b</sup> | 0.77* | 1.00 |  |  |
| BHS<sup>c</sup> | 0.63* | 0.64* | 1.00 |  |
| BAI<sup>d</sup> | 0.61* | 0.62* | 0.44* | 1.00 |

\* $p < .0001$

### Completion of treatment and Length of Stay (LOS)

Program completion was defined as satisfying all clinical requirements of the SACS program. Dropouts participated in the program but failed to complete.

### Completion rates

The overall completion rate was 23.5% (N=68). Significantly fewer higher MBP scoring patients completed the program (11.3%), than lower scoring patients (26.3%).

### Dropouts

N=221 (76.5%) met the criteria for dropout (i.e., self-termination of treatment without completing program requirements). Gender was not a factor for either completion or dropout (*X*^2^= 3.29; p=.07).

### Treatment Retention/Length of stay (LOS)

refers to the number of days that patients participated in treatment independent of whether they were completers or dropouts. The mean number of days in treatment for all patients (completers and dropouts) was 143.4 ± SD 7.55 with a range of 7-397 days. The median number of days spent in treatment was 100 days.

### Gender

Female patients participated in the program significantly longer that males (mean_females_ =163 days ± 14.1 vs mean _males_ = 132 days ± 8.6 days; t=1.98, df=287, p=.048).

### Completion Status

Completers stayed in treatment for an average of 197.4 days compared to 92.3 days for dropouts. This difference was highly significant (t=11.52, df=28, p<.0001). The earliest patient completed treatment in 94 days while the last patient required more than a year (397 days). Mean LOS with 95% Confidence Interval (CI) for all patients, completers, dropouts, high and low-moderate psychological pain intensity patient groups are illustrated in (Fig 1) for comparison.

**Fig 1.**
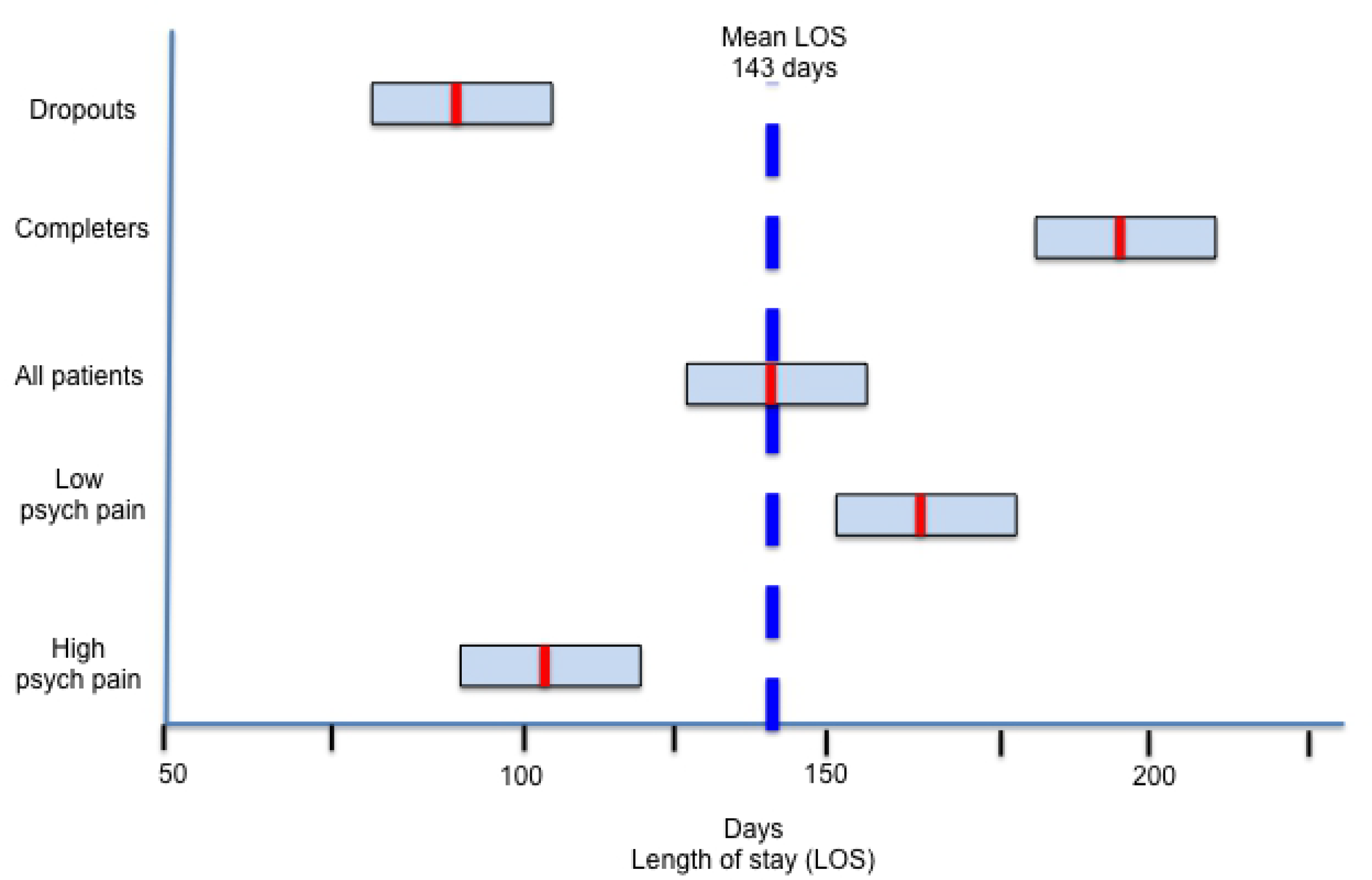
Mean retention times portrayed as LOS (days) with corresponding 95% CI for the total patient population, completers, dropouts, as well as high and low-moderate category psychological pain (MBP).

### Psychological Pain (MBP) ratings: Completers vs Dropouts

Although overall MBP scores were in the low moderate range for the total patient population (Table 2), psychological pain ratings were significantly higher for dropouts compared to completers (mean MBP_Dropouts_=24.4; mean MBP_Completers_= 20.9; t=-2.82, df=287, p=.005). MBP scores were also significantly higher in dropouts with briefer length of stays (LOS < 65 days) than for dropouts with longer LOS (mean MBP _<65LOS_ =25.9; mean MBP _>65 LOS_=23.1; t=5.24, df=219, p=.02).

### Correlation between length of stay (LOS) and ratings for psychological pain (MBP) depression (BDI), anxiety (BAI) and hopelessness (BHS)

Regression analysis revealed significant negative linear correlations between LOS and scores of all psychometric assessments with the exception of hopelessness (BHI). Psychological pain was most strongly correlated with length of stay (LOS) (r= −0.20, Cohen’s d=0.42, p=.001) compared with the other assessments. This was followed by depression (BDI r=-.165, Cohen’s d=0.33, p=.005) and anxiety (BDI r=-.135, Cohen’s d=0.27, p=.022). Hopelessness was non-significant (BHS r=-.107, Cohen’s d=0.22, p=.07).

### Relationship between Psychological Pain (MBP) and Clinical Outcomes

A subgroup of 53 patients (18.3%) meeting the criterion for scoring high on psychological pain assessment was identified. A threshold for high psychological pain (MBP ≥ 32) was developed and described in an earlier MBP scale validation study (10). As illustrated in Table 4, patients meeting this definition of high psychological pain scores (at intake) also rated significantly higher for depression (BDI), anxiety (BAI) and hopelessness (BHS) compared to lower MBP scoring patients.

**Table 4:**
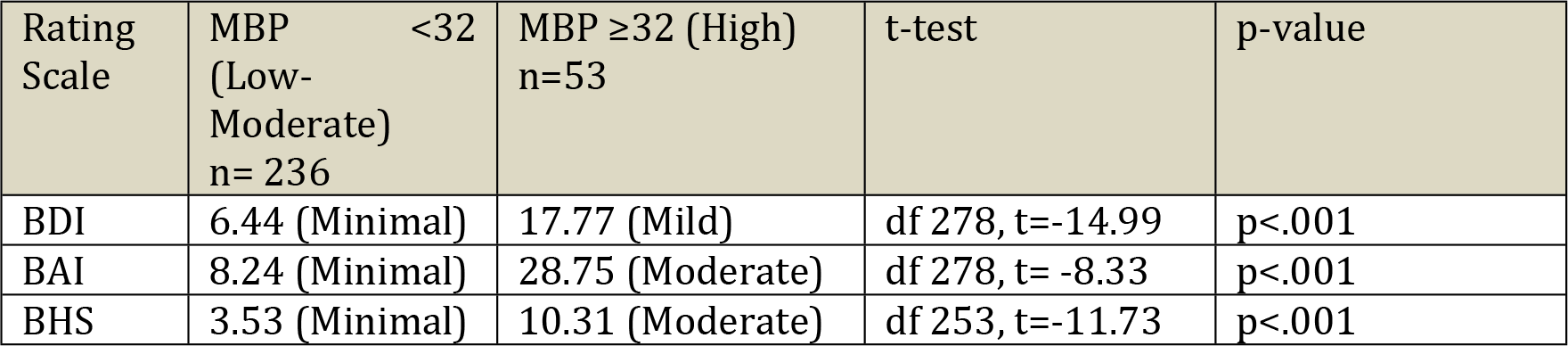
Significant differences in clinical assessment symptom severity between subgroups of patients scoring above and below threshold for high psychological pain (MBP)

Additionally, as shown in Table 5, higher scoring patients on MBP assessment had significantly diminished program retention times in terms of LOS and were more likely to drop out of treatment than the low-moderate group.

**Table 5.** Significant differences in clinical outcomes between patients scoring above and below-threshold for high psychological pain (MBP)

| Outcome measure | MBP $\geq 32$ (high) N=53 | MBP <32 (lower) N= 236 | p-value ($\chi^2$) |
| --- | --- | --- | --- |
| Completers | 6 (11.3%) | 62 (26.3%) | p<.02 |
| Dropouts | 47 (88.7%) | 174 (73.7%) | p=.02 |
| Length of Stay (Mean) | 102 days | 152 days | p<.009 |
| Dropout to Completion ratio | 7.8 | 2.79 | p<.02 |

### Completion Rates and LOS

Logistic regression analysis showed that patients with high category pre-treatment MBP scores were significantly more likely to become treatment dropouts compared to patients reporting lower intensity psychological pain (Odds ratio 2.79, RR 1.21, p=.025). Additionally, there were significantly more dropouts per completion in the high pain group (47:6,) compared to the lower scoring psychological pain group (174:62) [χ^2^ = 5.38, p=0.02].

Retention in treatment (LOS) was significantly reduced in high pain category patients relative to low-moderate patients (Table 5). A separate analysis replicated this pattern within the Dropout group where high-pain Dropout patients demonstrated a reduced LOS (mean=73.1d) relative to lower pain Dropouts (mean=97.5d) (p=.02, t=2.31, df =219). While high pain category patients exhibited diminished LOS and lower subsequent completion rates, we observed robust increases in both of these variables for patients who remained in treatment for more than 100 days. Specifically, completion rates for the high pain group increased from 11.3% to 35.3% and the low-moderate pain group increased from 26.3% to 48.3% when LOS was greater than 100 days. Overall, 96.8% of all completions for both groups occurred after 100 days of treatment.

Fig 1 illustrates mean LOS and 95% confidence interval (CI) for patients grouped by outcome status and psychological pain intensity at program admission. Notably, the mean LOS of high psychological pain (MBP) category patients was significantly decreased compared to the low-moderate pain patients.

### Retention Curves and Survival Analyses

Kaplan-Meier survival analyses and retention curves were developed for further comparison of the high (n=53) and low-moderate MBP (n=236) psychological pain subgroups for visualizing LOS and dropout patterns. These analyses revealed significant differences between the patient curves (Log Rank p=0.001) in terms of retention rates and patterns (Fig 2). Even at similar time points, the considerable over-representation of completions clustering on the Low-moderate pain curve while largely absent on the High pain curve is visually apparent. A separate analysis performed on the dropout group alone comparing the high and low-moderate pain groups reflected a similar difference between the curves (p=.011). 66% (n=35) of high scoring MBP patients had dropped out before the first patient completed the treatment program (day 94) and by day 129, 75% of high-pain patients had dropped out. High-pain category patients reached 50% attrition after just 53 days compared to 108 days for the lower pain group.

**Fig 2.**
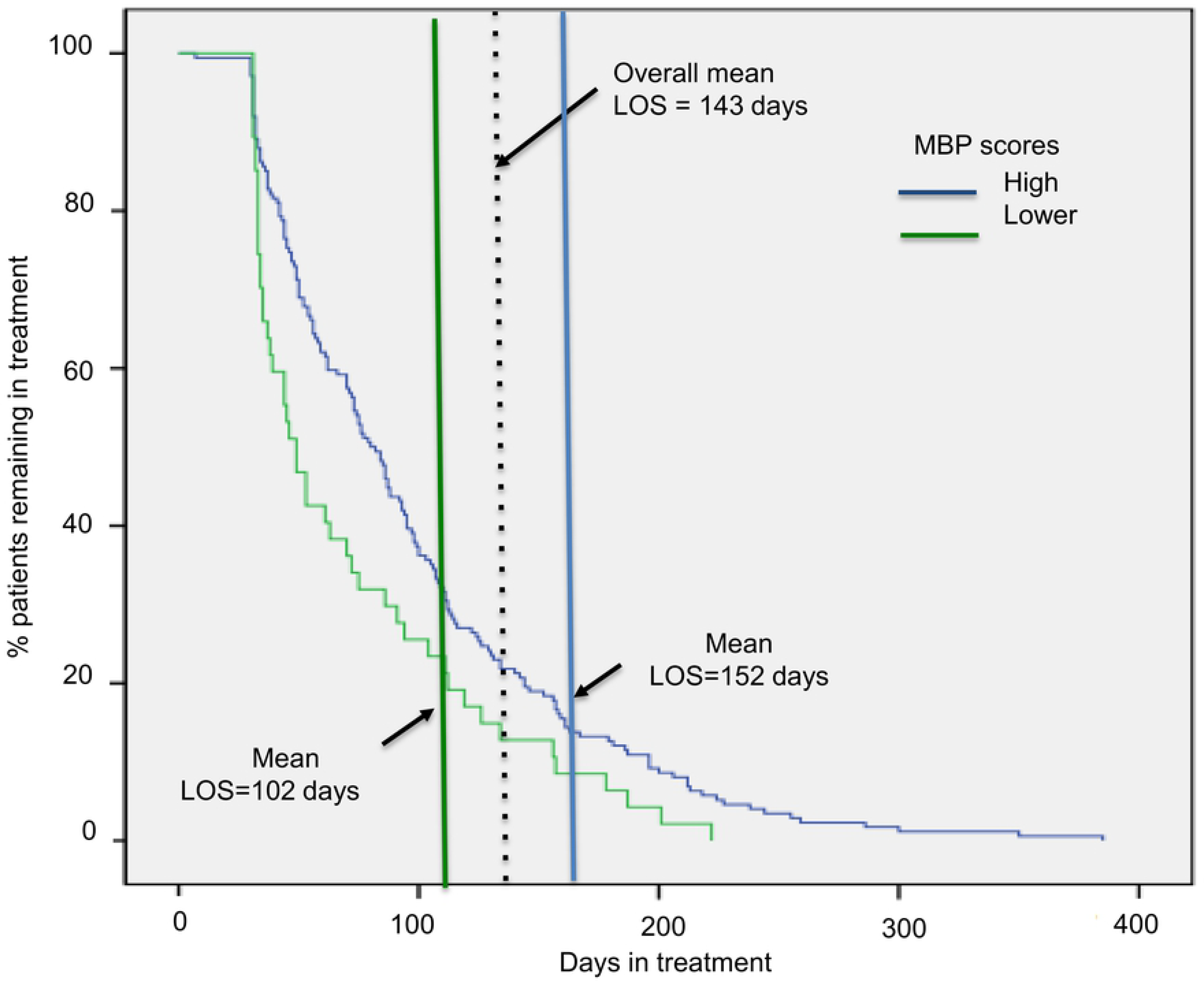
Program retention curves for High and Low-moderate pain categories. Log Rank analysis showed that the curves significantly differed between the high pain and low-moderate pain patient groups (χ2=11.1, p=.001). (+) indicates individual patient-program completion. There is a notable clustering of completions on the low-moderate pain curve while relatively absent on the high pain curve even at similar time points. At the mean LOS of 143d for the general group, nearly twice the percentage of low-moderate psychological pain patients remained in treatment compared with the high category pain patients (41.0% vs 22.5%).

## DISCUSSION

This study is, to our knowledge, the first to specifically focus on characterizing psychological pain in a population seeking treatment for substance use disorders. Primarily, the data from this effort confirm that psychological pain is a quantifiable construct in patients suffering from substance abuse/dependence and that the MBP demonstrated adequate reliability for measuring psychological pain in this clinical population. Additionally, we found evidence that elevated pretreatment psychological pain is associated with negative treatment outcomes such as diminished treatment retention time (LOS) and reduced likelihood for program completion. We chose program completion as a proximal indicator of overall treatment outcome although we did not have follow-up for abstinence.

Dropout was the most frequent treatment outcome for the SACS patients. This observation agrees with data from the Treatment Episode Data Set (TEDS) published by SAMHSA (32). Our patient population exhibited dropout rates somewhat higher than many programs reported to the TEDS nationwide. It is possible that this was in part due to a high proportion (73.7%) of our patients reporting methamphetamine dependence. Data from a similar region in Los Angeles, California found that methamphetamine abusers were likely to continue drug use during initial entry into the program, a factor which was thought to contribute to the relatively high attrition rate (51%) during the first few weeks of treatment (33). Another potential factor is that many of the patients were disadvantaged. We did find that completion rates in the SACS population nearly doubled when patients remained in treatment at least 100 days, reaching levels in agreement with data from the TEDS Government analysis.

Pre-treatment levels of anxiety, depression and hopelessness symptoms for the entire treatment population were indicative of minimal comorbid psychopathology. A simple binary risk stratification method based on psychological pain (MBP scores), previously developed and applied to depressed and suicidal psychiatric populations (9, 10) re-grouped patient data into high and low-moderate categories of pain. The ‘high’ pain category group, relative to the lower pain group, exhibited significantly greater dropout rates, had more severe psychopathology (depression and anxiety) scores as well as a pronounced reduction in LOS. Survival curve analyses confirmed differences in completion patterns and LOS which suggest that our high and low-moderate pain risk categorization scoring method separated patients into two sub-populations differing in treatment outcomes. Incorporating systematic psychological pain screening within current standard intake assessment paradigms, may aid in identifying patients at program entry posing elevated risk for early dropout and offer the potential for outcome modifying interventions such as increasing retention time. For example, each of the relatively few high-pain category patients who successfully completed the treatment protocol (only 2.1% of the total patient sample) were associated with LOS >129 days; nearly twice the mean LOS for the total high pain population. In contrast, 89% of high pain patients who dropped out of treatment, did so before the first 102 days of treatment.

This is a small, retrospective observational study. In light of the preliminary nature of these findings, caution is warranted in generalizing them pending replication in larger populations. Future replicative studies would benefit from a prospective design, however, retrospective designs can be appropriately used in the context of multiple outcome measures (34). Demographic information was limited to and dependent upon the clinical program intake process. An additional limitation was that these findings derive from a relatively small sample size. Statistical significance, however, was reached for nearly all tests performed, reflecting the robust effect size we have observed in our previous studies on the effects of elevated psychological pain on clinical outcomes.

## CONCLUSION

In this study, we present evidence suggesting that psychological pain has a negative influential effect on program completion and LOS in outpatient substance treatment. The highest scoring patients on pre-treatment psychological pain assessment were ultimately 1.21 times more likely to drop out and to participate in treatment significantly fewer days compared with lower pain scoring peers. Whether this reduction in completion rates is a direct or indirect consequence of decreased LOS remains unanswered and further work in larger populations is needed to better understand these relationships. The survival curve analysis demonstrated a preferential clustering of completions on the lower pain group curve and relative lack of completions on the higher pain curve at identical time points. This suggests that a factor apart from reduced LOS may also be negatively influencing completion likelihood. Regardless, the study of psychological pain represents a novel area to further our understanding of the unpredictable outcomes in substance use disorders treatment. The subset of patients experiencing very high levels of psychological pain at treatment initiation may be inherently poorer candidates for outpatient substance treatment and early identification could allow for prompt referral to accessing higher levels of care. Efforts to further our understanding on the negative influence of high pre-treatment levels of psychological pain on completion rates and LOS offer additional opportunities for improving substance treatment outcomes.

